# Janus kinase 2 regulates Nurr1 protein stability in dopaminergic neurons of the aging midbrain

**DOI:** 10.64898/2026.03.19.712974

**Authors:** Yongwoo Jang, Young Hwa Kim, Jeha Jeon, Young Cha, Claudia Lopes, Jin Hyuk Jung, Eunji Oh, Yeongwon Park, Chanyoung Ko, Bobae Hyeon, Pierre Leblanc, Kwang-Soo Kim

## Abstract

The nuclear receptor Nurr1 (NR4A2) is an essential transcription factor that governs the differentiation, maturation, and long-term maintenance of midbrain dopaminergic (mDA) neurons in the substantia nigra. Reduced Nurr1 expression has been closely linked to age-related dopaminergic neuronal loss and the pathogenesis of Parkinson’s disease. However, the molecular mechanisms regulating Nurr1 expression and protein stability in the aging midbrain remain poorly understood. Here, we identify Janus kinase 2 (JAK2) as a previously unrecognized regulator of Nurr1 in mDA neurons. In the substantia nigra of aged mice (12-and 18-month-old), JAK2 was robustly expressed in Nurr1-positive mDA neurons, whereas its expression was minimal in young adult mice. In SK-N-BE(2)C neuroblastoma cells, overexpression of JAK2 modestly enhanced Nurr1 transcriptional activity, while the constitutively active mutant JAK2_V617F_ markedly increased it. Notably, this effect was not blocked by pharmacological inhibition of STAT, PI3K, or Akt signaling pathways, indicating that JAK2 regulates Nurr1 independently of canonical JAK/STAT or PI3K/Akt signaling. Mechanistically, JAK2 did not promote tyrosine phosphorylation of Nurr1 but instead physically interacted with Nurr1, leading to enhanced nuclear stability of the Nurr1 protein. Consistent with this mechanism, expression of JAK2_V617F_ increased Nurr1 protein levels without altering its mRNA expression. Functionally, co-expression of JAK2_V617F_ and Nurr1 attenuated oxidative stress–induced cytotoxicity and reduced reactive oxygen species accumulation. Together, these findings reveal a phosphorylation-independent mechanism by which JAK2 stabilizes Nurr1 protein and enhances its transcriptional activity. Our results further suggest that age-associated induction of JAK2 in dopaminergic neurons may promote neuronal resilience by maintaining Nurr1 protein stability during aging.

## INTRODUCTION

Midbrain dopaminergic (mDA) neurons release the neurotransmitter dopamine and regulate motor function through the projections from the substantia nigra pars compacta (SNpc) to the dorsal striatum, forming the nigrostriatal pathway ^1^. The long-term stability and precise regulation of these neurons are therefore essential for maintaining normal motor function. Degeneration of mDA neurons in the SNpc represents a central pathological hallmark of Parkinson’s disease (PD), an age-associated neurodegenerative disorder characterized by progressive motor dysfunction ^2, 3^.

The nuclear receptor Nurr1 (NR4A2) is a key transcription factor required for the development, long-term maintenance, and protection of mDA neurons from inflammation-induced degeneration ^4–6^. Nurr1 expression begins during dopaminergic lineage specification and persists throughout adulthood in mature dopaminergic neurons ^7^. In these neurons, Nurr1 regulates the transcription of genes essential for dopamine synthesis, reuptake, and storage, including tyrosine hydroxylase (TH), aromatic L-amino acid decarboxylase (AADC), dopamine transporter (DAT), and vesicular monoamine transporter 2 (VMAT2) ^4, 8, 9^. Consistent with this role, Nurr1-deficient mice fail to generate mDA neurons ^5^, whereas conditional disruption of Nurr1 in adulthood leads to progressive loss of striatal dopaminergic terminals ^10^. Collectively, these findings establish Nurr1 as an indispensable regulator of the differentiation and long-term integrity of mDA neurons.

Reduced Nurr1 expression has been reported in aging human brains and in patients with PD, as well as in experimental models involving aging, α-synuclein overexpression, and dopaminergic neurotoxins ^11–15^. These observations suggest that impaired Nurr1 expression or signaling contributes to dopaminergic neuronal vulnerability during aging and PD pathogenesis. However, the molecular mechanisms regulating Nurr1 expression and stability in the aging midbrain remain largely unknown.

Aging of the midbrain is accompanied by increased oxidative stress and altered inflammatory signaling, both of which contribute to dopaminergic neuronal vulnerability. The concept of *inflammaging* highlights the interplay between chronic inflammation and aging in neurodegenerative processes. Cytokine- and stress-responsive signaling pathways may therefore influence transcriptional mechanisms that maintain dopaminergic neuron homeostasis. Among such pathways, Janus kinase 2 (JAK2) is a central mediator of cytokine and stress signaling and serves as an important interface between immune signaling and cellular responses ^16^. JAK2 is a cytoplasmic tyrosine kinase best characterized in immune and hematopoietic systems, where it is activated downstream of cytokine and growth factor receptors ^17, 18^. A constitutively active mutant, JAK2_V617F_, identified in myeloproliferative disorders, confers ligand-independent constitutive activation of JAK2 and has been extensively used to investigate JAK2 signaling mechanisms ^19–21^. Activated JAK2 typically signals through the JAK/STAT and phosphatidylinositol 3-kinase (PI3K)/Akt pathways ^22–25^.

In the central nervous system, JAK1 and JAK2 are highly expressed during embryonic development, with expression declining into adulthood ^26^. JAK2/STAT signaling has also been implicated in neuronal survival in models of neurodegeneration, including protection against amyloid-β-induced toxicity in Alzheimer’s disease models ^23, 27–30^. However, the expression and functional significance of JAK2 in mDA neurons remain largely unexplored.

Given the central role of Nurr1 in mDA neuron survival and its involvement in neuroinflammatory processes, we hypothesized that age-associated signaling pathways may involve JAK2 in the regulation of Nurr1. In the present study, we examined the expression pattern of JAK2 in mDA neurons during aging and investigated its potential role in regulating Nurr1 expression and function in these neurons.

## RESULTS

### Age-related expression of JAK2 in Nurr1-positive dopaminergic neurons

To assess the potential relevance of JAK2 in the midbrain, we first examined its expression in midbrain dopaminergic (mDA) neurons of the mouse brain. In 2-month-old mice, JAK2 immunoreactivity was barely detectable in the substantia nigra (SN) (Figure 1A). Because age-dependent dopaminergic neuronal loss is a hallmark of Parkinson’s disease (PD), we next analyzed JAK2 expression across aging stages (2, 6, 12, and 18 months). Surprisingly, JAK2 immunofluorescence intensity increased progressively with age. In 12-and 18-month-old mice, JAK2 expression was prominently observed in tyrosine hydroxylase (TH)-positive dopaminergic neurons of the substantia nigra (Figure 1B,C). Quantitative analysis further demonstrated that the proportion of JAK2-positive cells among total TH-positive neurons increased in an age-dependent manner (Figure 1D).

**Figure 1.**
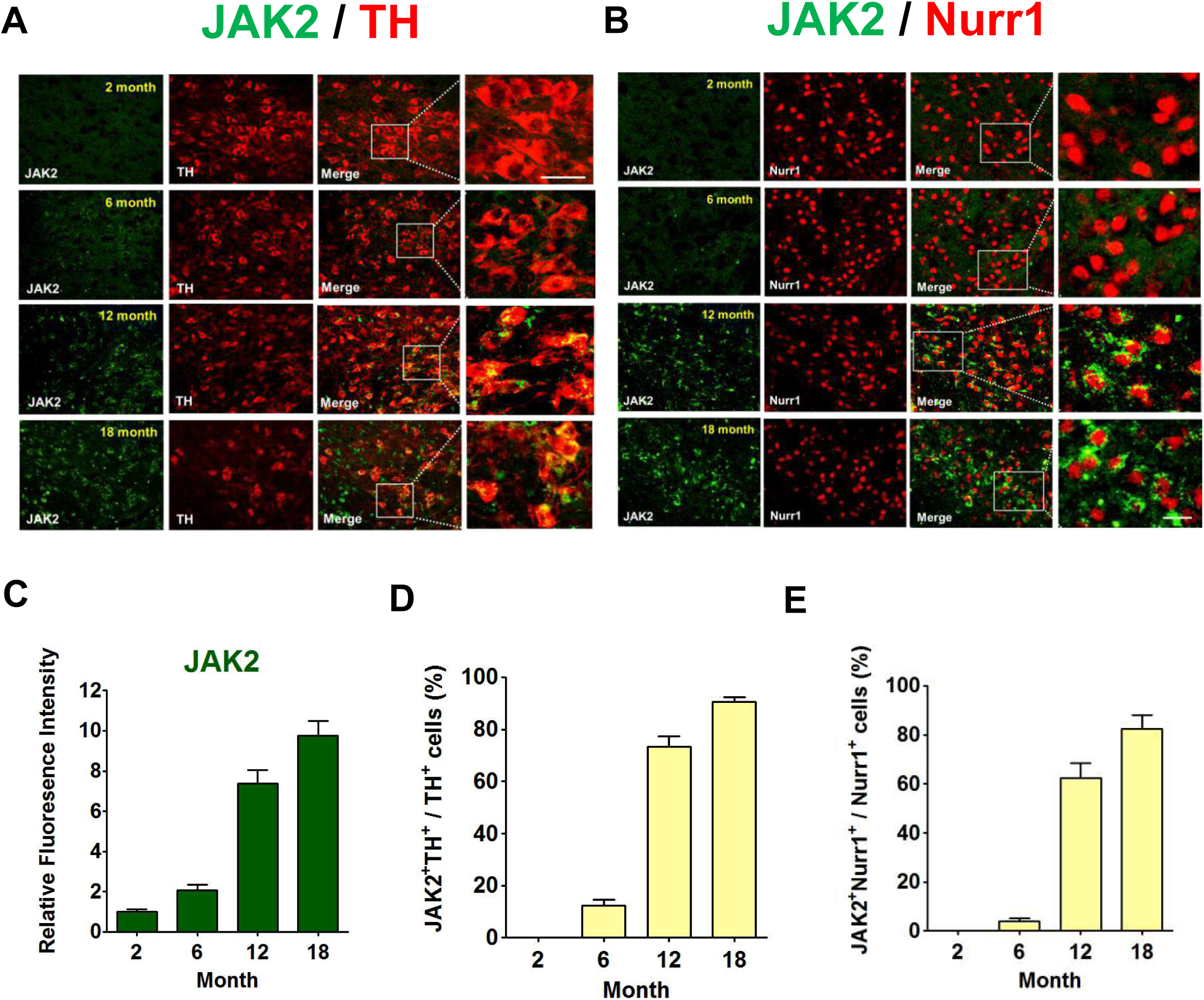
Age-dependent JAK2 expression in midbrain dopaminergic neurons (A-B) Representative immunofluorescence images showing JAK2 expression in the substantia nigra pars compacta (SNc) of mice at 2, 6, 12, and 18 months of age. Sections were co-stained with antibodies against JAK2 and either tyrosine hydroxylase (TH) (A) or Nurr1 (B). JAK2 immunoreactivity was barely detectable in 2-month-old mice but became evident from 6 months of age and increased further in 12- and 18-month-old mice. Scale bar: 50 μm. (C) Quantification of relative JAK2 fluorescence intensity in the SNc at each age after normalization to background fluorescence. (D-E) Quantification of the percentage of JAK2-positive cells among TH-positive neurons (D) and Nurr1-positive neurons (E) across age groups. Data are presented as mean ± SEM.

Previous studies have reported that reduced Nurr1 expression is closely associated with aging and degeneration of dopaminergic neurons. We therefore compared the expression patterns of JAK2 and Nurr1 in the midbrain across aging stages. As shown in Figure 1B, age-associated JAK2 expression was largely co-localized with Nurr1-positive dopaminergic neurons. Quantitative analysis revealed that the proportion of JAK2-positive cells among total Nurr1-positive neurons also increased with age (Figure 1E). Interestingly, JAK2 expression was not induced in degenerating dopaminergic neurons following stereotaxic injection of the dopaminergic neurotoxin 6-hydroxydopamine (6-OHDA) into the substantia nigra (Figure S1). Together, these findings indicate that JAK2 upregulation in dopaminergic neurons occurs in an aging-associated manner rather than as a general response to neurotoxic injury.

### JAK2-mediated activation of Nurr1 transcriptional activity

To determine whether increased JAK2 expression affects Nurr1 function, we examined Nurr1 transcriptional activity following JAK2 overexpression. Luciferase reporter assays were performed in SK-N-BE(2)C neuroblastoma cells using reporter constructs containing either an NGFI-B responsive element (NBRE)-like motif (NL3), which serves as a monomeric Nurr1 binding site, or a direct repeat 5 (DR5) motif, which mediates heterodimeric binding of Nurr1 with RXR. As shown in Figure 2A, overexpression of wild-type (WT) JAK2 modestly increased Nurr1 transcriptional activity in the NL3 reporter assay (blue bar). In contrast, NL3-driven Nurr1 transcriptional activity was markedly enhanced by overexpression of the constitutively active JAK2_V617F_ mutant, a gain-of-function mutation frequently identified in patients with myeloproliferative neoplasms ^20, 31^.

**Figure 2.**
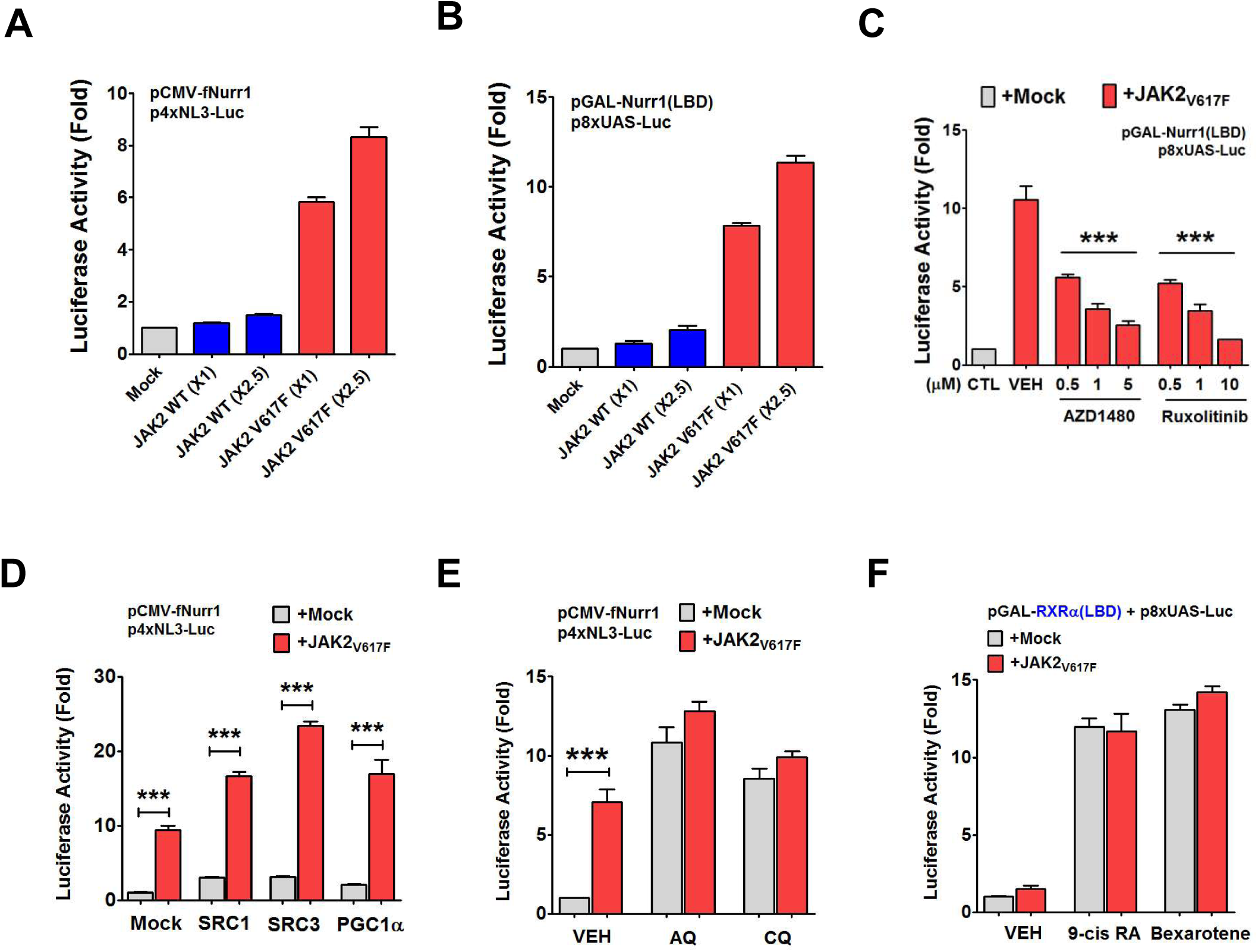
JAK2_V617F_-induced Nurr1 transcriptional activation (A-B) Luciferase reporter assays showing the effect of JAK2_V617F_, a constitutively active mutant, on the transcriptional activity of full-length Nurr1 (pCMV-fNurr1) (A) and GAL4-Nurr1(LBD) fusion protein (B) in SK-N-BE(2)C cells. (C) JAK2_V617F_-induced Nurr1 activation was significantly suppressed by pharmacological JAK2 inhibitors, AZD1480 and ruxolitinib. *p < 0.05, **p < 0.01, ***p < 0.001; one-way ANOVA followed by Tukey’s *post hoc* test. (D) Co-expression of transcriptional coactivators (SRC1, SRC3, and PGC1α) further potentiated JAK2_V617F_-induced Nurr1 transcriptional activity in SK-N-BE(2)C cells. ***p < 0.001; Student’s *t*-test. (E) JAK2_V617F_-induced Nurr1 activation was not affected by Nurr1 agonists, amodiaquine (AQ) or chloroquine (CQ). (F) JAK2_V617F_ did not alter the transcriptional activity of RXRα, the heterodimeric partner of Nurr1.

We next assessed whether JAK2 influences Nurr1–RXRα heterodimer–mediated transcription using the DR5 reporter in the presence of the RXRα agonists 9-cis retinoic acid or bexarotene. Neither WT JAK2 nor JAK2_V617F_ altered DR5-driven Nurr1 transcriptional activity (Figure S2). To further determine whether JAK2_V617F_ acts on the ligand-binding domain (LBD) of Nurr1, we employed an additional reporter construct in which the GAL4 DNA-binding domain was fused to the Nurr1 LBD. Consistently, JAK2_V617F_ significantly increased Nurr1-LBD–dependent transcriptional activity (Figure 2B), suggesting that active JAK2 modulates the transcriptional activity of Nurr1. Importantly, JAK2_V617F_-induced activation of Nurr1 was abolished by pharmacological inhibition of JAK2 using AZD1480 and ruxolitinib ^32, 33^, indicating that Nurr1 activation is dependent on JAK2 kinase activity (Figure 2C).

We next examined whether Nurr1 coactivators (SRC1, SRC3, and PGC1α) modulate JAK2_V617F_-induced Nurr1 transcriptional activity. As previously reported ^34^, overexpression of SRC1, SRC3, or PGC1α enhanced Nurr1 transcriptional activity (Figure 2D, grey bars). Co-expression of JAK2_V617F_ further increased Nurr1 transcriptional activity in the presence of these coactivators (Figure 2D, red bars). In contrast, JAK2_V617F_ did not alter Nurr1 transcriptional activation induced by the Nurr1 agonists amodiaquine (AQ) or chloroquine (CQ) ^35, 36^ (Figure 2E). Because RXRα functions as a heterodimeric partner of Nurr1, we further examined whether JAK2 affects RXRα transcriptional activity. As shown in Figure 2F, JAK2 did not influence RXRα-LBD–dependent transcription in either the absence or presence of RXRα agonists.

To determine whether JAK2 selectively activates Nurr1, we examined the effect of JAK2_V617F_ on the transcriptional activity of other nuclear receptors. Among NR4A family members, JAK2_V617F_ enhanced Nor1 (NR4A3) transcriptional activity, although to a lesser extent than Nurr1, while no significant effect was observed for Nur77 (NR4A1) (Figure 3A). We next assessed whether the effect of JAK2_V617F_ extends to other representative nuclear receptors, including peroxisome proliferator-activated receptor α (PPARα), PPARγ, glucocorticoid receptor (GR), and liver X receptor (LXR). Using GAL4-based luciferase reporter assays, we confirmed that each receptor was functionally activated by its respective ligand (Figure 3). However, JAK2_V617F_ did not affect the transcriptional activity of PPARα-LBD (Figure 3B), PPARγ-LBD (Figure 3C), GR-LBD (Figure 3D), or LXR-LBD (Figure 3E). Collectively, these results indicate that JAK2_V617F_ selectively enhances the transcriptional activity of Nurr1 and, to a lesser extent, Nor1.

**Figure 3.**
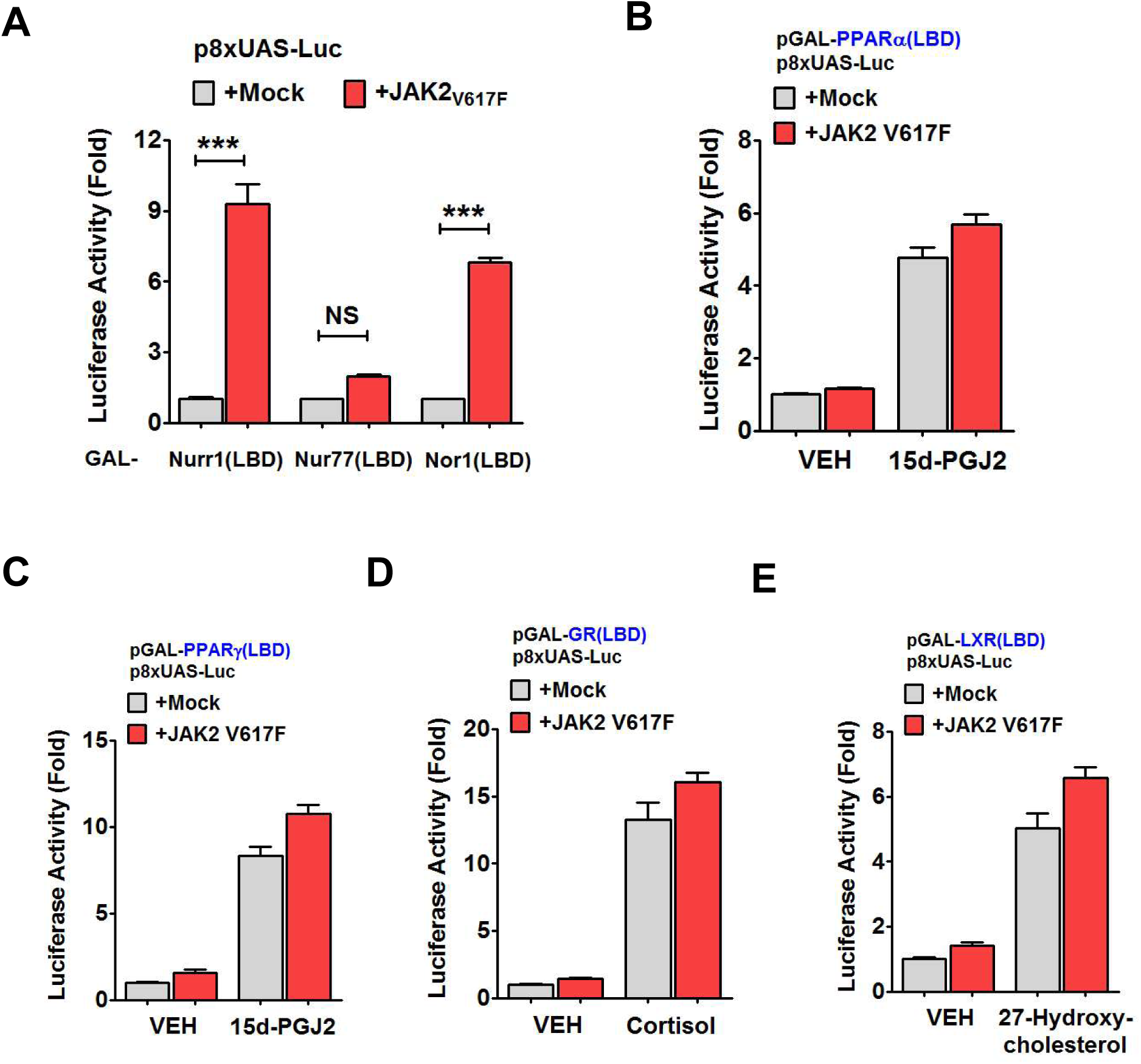
Effect of JAK_V617F_ on transcriptional activity of other nuclear receptors (A) Luciferase reporter assay showing that JAK2_V617F_ significantly enhanced Nor1 transcriptional activity, whereas no effect was observed on Nur77. ***p < 0.001, Student’s *t* test (B-C) JAK2_V617F_ did not alter ligand-induced transcriptional activity of PPARα (B) or PPARγ (C) in the presence of 15d-PGJ2. (D-E) JAK2_V617F_ had no effect on ligand-induced transcriptional activity of the glucocorticoid receptor (GR) activated by cortisol (D) or liver X receptor (LXR) activated by 27-hydroxycholesterol (E).

### JAK2–Nurr1 interaction independent of canonical JAK/STAT and PI3K/Akt signaling

JAK2 is well known for its association with STAT transcription factors as part of the canonical JAK–STAT signaling pathway ^37^. We therefore examined whether STAT proteins contribute to JAK2_V617F_-induced Nurr1 transactivation. In the GAL4–Nurr1-LBD luciferase reporter system, overexpression of STAT3 did not alter JAK2_V617F_-induced Nurr1 transcriptional activity (Figure S3A). Consistently, pharmacological inhibition of STAT signaling using fludarabine (STAT1 inhibitor), S3I-201 (STAT3 inhibitor), or a STAT5 inhibitor did not suppress JAK2_V617F_-mediated Nurr1 activation (Figure S3B).

We next assessed the involvement of the phosphoinositide 3-kinase (PI3K)/Akt pathway, another signaling cascade regulated by JAK2^24, 25^. Inhibition of PI3K with wortmannin or LY294002, as well as inhibition of Akt with GSK690693 or MK-2206, failed to attenuate JAK2_V617F_-induced Nurr1 transcriptional activity (Figure S4). Together, these findings indicate that JAK2_V617F_-mediated activation of Nurr1 occurs independently of canonical JAK/STAT and PI3K/Akt signaling pathways.

Because JAK2 functions as a tyrosine kinase, we next examined whether JAK2 directly phosphorylates Nurr1. Based on the results shown in Figure 2B, which indicated that the LBD of Nurr1 is required for JAK2_V617F_-mediated activation, we analyzed the tyrosine residues within the Nurr1-LBD. To assess the potential contribution of these residues, we generated Nurr1-LBD mutants in which individual tyrosine (Y) residues (Y384, Y393, Y407, Y453, Y551, and Y575) were substituted with phenylalanine (F), which lacks the hydroxyl group required for phosphorylation. As shown in Figure 4A, JAK2V617F retained the ability to enhance the transcriptional activity of each Nurr1 mutant. Because multiple tyrosine residues could potentially contribute to phosphorylation, we next predicted candidate phosphorylation sites within the Nurr1 sequence using the Group-based Prediction System (Ver.3, http://gps.biocuckoo.org) (Figure S5). Among the tyrosine residues within the Nurr1-LBD, Y384, Y393, and Y575 were predicted as potential JAK2 phosphorylation sites (Figure S5). To evaluate their contribution, we generated Nurr1 mutants containing combinatorial substitutions at these residues, as illustrated in Figure 4B. However, none of these mutants exhibited altered responsiveness to JAK2_V617F_ in the transcriptional assay (Figure 4B).

**Figure 4.**
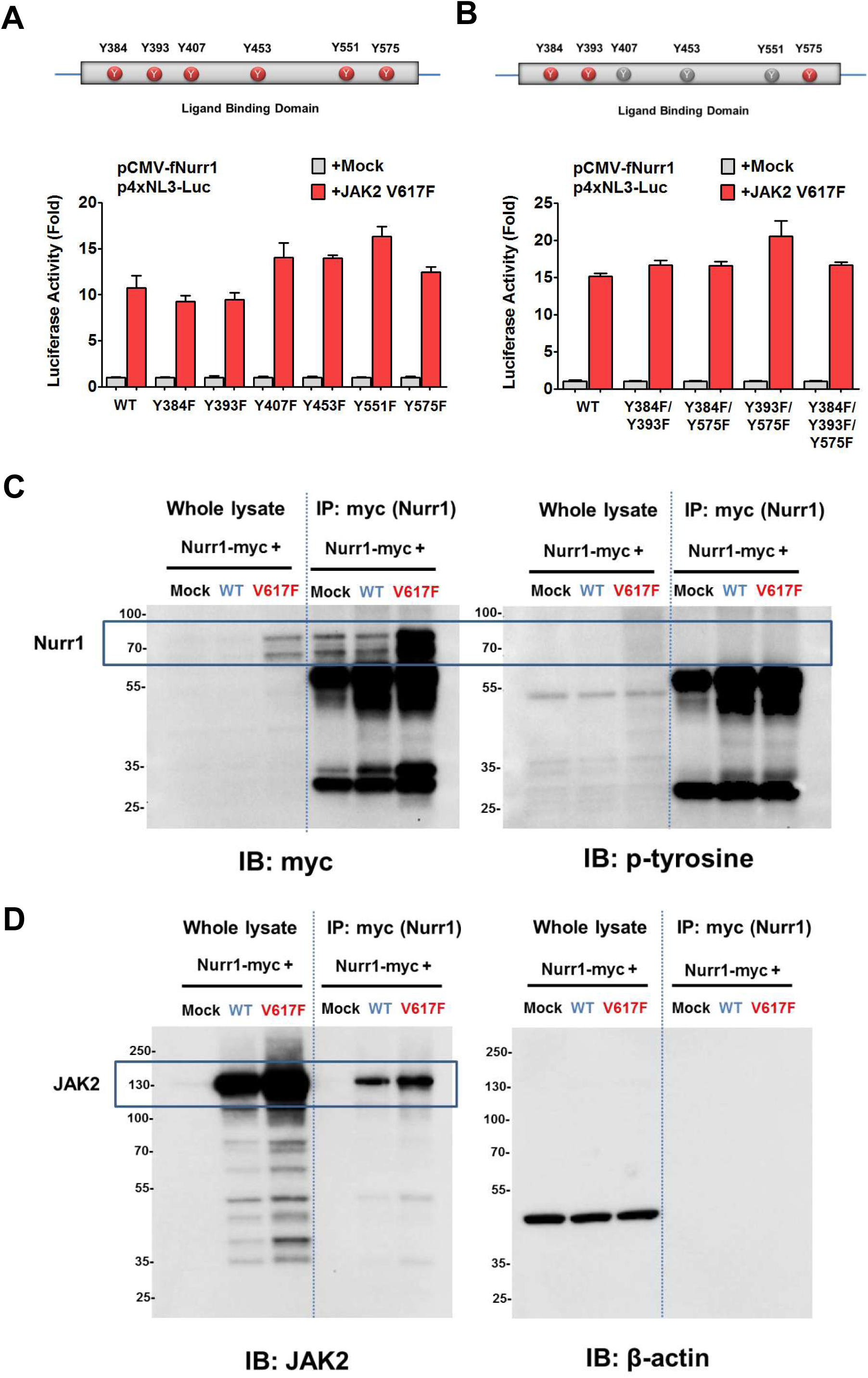
Phosphorylation-independent interaction of JAK2 with Nurr1-LBD (A) Luciferase reporter assays showing that single tyrosine (Y)-to-phenylalanine (F) substitutions within the Nurr1-LBD did not abolish JAK2_V617F_-induced transcriptional activation. (B) Double and triple Y-to-F mutations at putative JAK2 recognition sites predicted by a group-based phosphorylation prediction system also retained responsiveness to JAK2_V617F_. (C) Immunoblot analysis using a pan–phospho-tyrosine antibody revealed no significant difference in Nurr1 tyrosine phosphorylation levels in the presence or absence of JAK2_V617F_. (D) Co-immunoprecipitation analysis showing detection of JAK2 in Nurr1-LBD–associated protein complexes, indicating a physical interaction between JAK2 and Nurr1-LBD.

To further determine whether JAK2 directly phosphorylates Nurr1-LBD, we examined tyrosine phosphorylation levels of Nurr1 following JAK2 overexpression. Nurr1-myc was immunoprecipitated using an anti-myc antibody, and tyrosine phosphorylation was analyzed by immunoblotting with a pan–phospho-tyrosine antibody. No significant difference in Nurr1 tyrosine phosphorylation was detected in the presence or absence of JAK2 (Figure 4C), suggesting that JAK2 regulates Nurr1 transcriptional activity independently of direct tyrosine phosphorylation. Interestingly, immunoblot analysis of Nurr1 immunoprecipitates revealed co-precipitation of JAK2, supporting a biochemical interaction between JAK2 and Nurr1 (Figure 4D). Detection of JAK2 in the Nurr1-bound fraction indicates a physical association between JAK2 and Nurr1 under the experimental conditions.

### JAK2-mediated stabilization of Nurr1 protein

Because the protein stability of nuclear receptors is closely linked to their transcriptional activity ^38^, we examined whether JAK2 influences Nurr1 expression at the mRNA and protein levels in SK-N-BE(2)C cells. Co-overexpression experiments revealed that Nurr1 mRNA levels were not altered by either wild-type (WT) JAK2 or JAK2_V617F_ (Figure 5A). In contrast, Nurr1 protein levels were markedly increased upon JAK2_V617F_ overexpression (Figure 5B). The absence of changes in Nurr1 mRNA expression suggest that the increase in protein levels results from post-transcriptional regulation, potentially through enhanced protein stability. Consistent with this interpretation, JAK2_V617F_ overexpression produced a pronounced increase in nuclear Nurr1-GFP fluorescence intensity (Figure 5C), indicating increased accumulation of Nurr1 protein in the nucleus. To directly evaluate Nurr1 protein stability, we performed cycloheximide chase assays to inhibit de novo protein synthesis. As shown in Figure 5D, cycloheximide treatment resulted in a time-dependent decrease in Nurr1 protein levels. Notably, overexpression of JAK2_V617F_ significantly attenuated this degradation, maintaining higher Nurr1 protein levels compared with control cells. Together, these results indicate that JAK2_V617F_ enhances the stability of Nurr1 protein.

**Figure 5.**
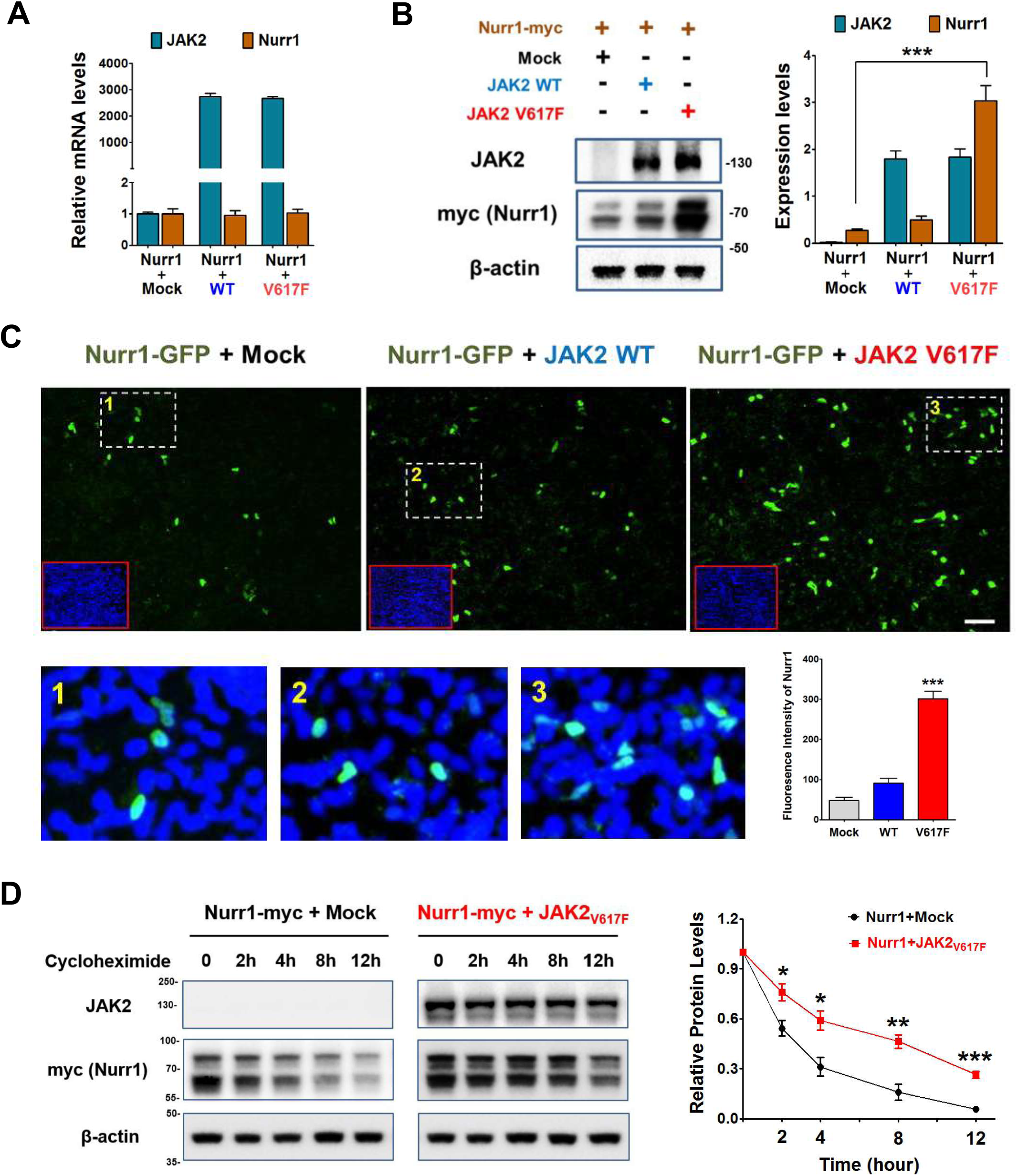
JAK2-mediated stabilization of Nurr1 protein (A) Quantitative real-time PCR analysis showing relative Nurr1 mRNA levels following overexpression of mock vector, wild-type (WT) JAK2, or JAK2V617F. No significant differences in Nurr1 mRNA levels were observed among groups. (B) Immunoblot analysis and quantification of Nurr1 protein levels following overexpression of mock vector, WT JAK2, or JAK2_V617F_. JAK2_V617F_ significantly increased Nurr1 protein levels. ***p < 0.001, Student’s *t*-test (C) Representative fluorescence images and quantification of nuclear Nurr1-GFP intensity in SK-N-BE(2)C cells. JAK2_V617F_ significantly increased nuclear Nurr1-GFP fluorescence. ***p < 0.001; one-way ANOVA with Tukey’s *post hoc* test (D) Cycloheximide chase assay showing that JAK2_V617F_ delayed Nurr1 protein degradation compared with mock-transfected cells. *p < 0.05, **p < 0.01, ***p < 0.001; Student’s *t*-test

### Protective effects of JAK2_V617F_ and Nurr1 against oxidative stress

In the aged brain, increased oxidative stress is a hallmark of age-related neuronal degeneration ^39^. Given the established role of Nurr1 in protecting against cellular damage, including reactive oxygen species (ROS)–mediated stress, we examined the effects of JAK2_V617F_ and Nurr1 under oxidative conditions. Exposure of SK-N-BE(2)C cells to hydrogen peroxide (H₂O₂) induced cytotoxicity in a concentration-dependent manner (Figure 6A, left). Overexpression of either JAK2_V617F_ or Nurr1 significantly reduced H₂O₂-induced cytotoxicity, whereas co-overexpression of JAK2_V617F_ and Nurr1 produced a greater protective effect (Figure 6A, right). A similar protective pattern was observed in rotenone-induced oxidative stress, a model of mitochondrial dysfunction (Figure 6B).

**Figure 6.**
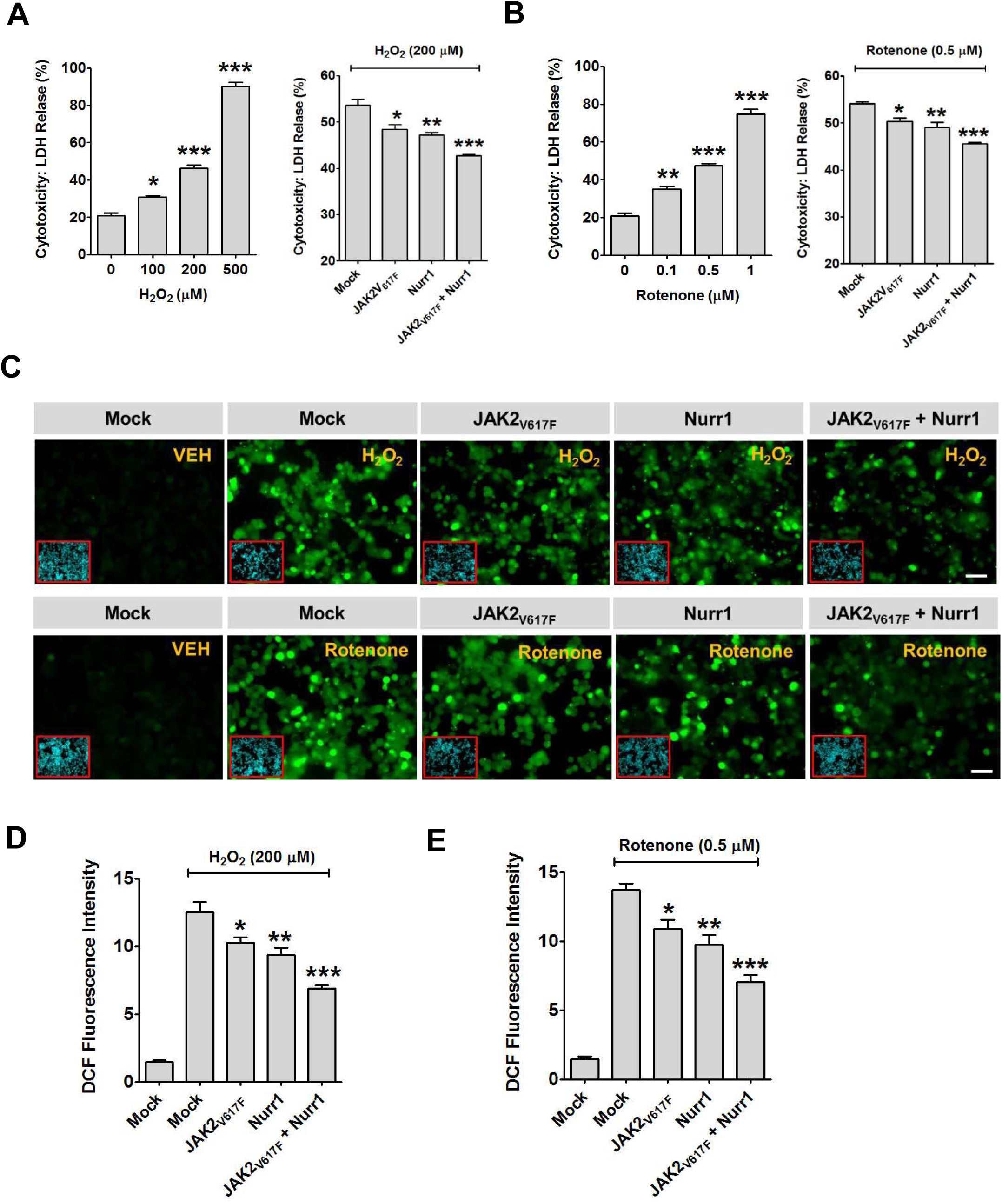
Protective effects of JAK2V617F and Nurr1 against oxidative stress–induced cytotoxicity and ROS accumulation (A) Hydrogen peroxide (H₂O₂) induced cytotoxicity in a concentration-dependent manner in SK-N-BE(2)C cells. Co-overexpression of JAK2_V617F_ and Nurr1 significantly attenuated H₂O₂-induced cytotoxicity. *p < 0.05, **p < 0.01, ***p < 0.001; one-way ANOVA with Tukey’s *post hoc* test (B) Rotenone induced cytotoxicity in a concentration-dependent manner in SK-N-BE(2)C cells. Co-overexpression of JAK2_V617F_ and Nurr1 significantly attenuated rotenone-induced cytotoxicity. *p < 0.05, **p < 0.01, ***p < 0.001, one-way ANOVA, Tukey’s *post hoc* test (C) Representative fluorescence images and quantification of intracellular reactive oxygen species (ROS) levels measured using 2′,7′-dichlorofluorescein diacetate (DCFH-DA) following exposure to H₂O₂ or rotenone in mock-, JAK2_V617F_-, Nurr1-, or JAK2_V617F_+ Nurr1–overexpressing cells. (D-E) Quantification of DCF fluorescence intensity showing that overexpression of JAK2_V617F_ or Nurr1 reduced H₂O₂-induced (D) and rotenone-induced (E) ROS accumulation. Co-expression of JAK2_V617F_ and Nurr1 produced an additional protective effect. *p < 0.05, **p < 0.01, ***p < 0.001; one-way ANOVA with Tukey’s *post hoc* test.

To quantify intracellular ROS levels, we used the fluorescent probe 2′,7′-dichlorofluorescin diacetate (DCFH-DA), which emits green fluorescence upon oxidation by ROS. Treatment with H₂O₂ or rotenone markedly increased DCF fluorescence in control cells. In contrast, overexpression of JAK2_V617F_ or Nurr1 significantly reduced ROS accumulation, with co-expression producing a further reduction in fluorescence intensity (Figure 6C and 6E). Collectively, these findings demonstrate that JAK2_V617F_ and Nurr1 cooperatively mitigate oxidative stress–induced cytotoxicity and ROS accumulation.

## ACKNOWLEDGMENTS

This work was in part supported by NIH grants (NS129188 and NS127391) and the Korea-US Collaborative Research Project (RS-2024-00468036).

## DISCUSSION

The nuclear receptor Nurr1 is a critical transcriptional regulator required for the development and long-term maintenance of midbrain dopaminergic (mDA) neurons ^40^. Numerous studies have demonstrated that reduced Nurr1 expression is closely associated with degeneration of mDA neurons in the aging brain and in patients with Parkinson’s disease (PD). However, the molecular mechanisms regulating Nurr1 expression and protein stability in aging mDA neurons remain poorly understood. In the present study, we identify JAK2 as a previously unrecognized regulator of Nurr1 and demonstrate that JAK2 enhances Nurr1 protein stability through a mechanism independent of canonical JAK/STAT and PI3K/Akt signaling pathways. These findings reveal a noncanonical mode of JAK2 signaling that directly modulates the stability and transcriptional activity of a key regulator of dopaminergic neuron maintenance.

Age-related degeneration of mDA neurons is strongly associated with late-onset PD ^41^. Post-mortem analyses of human brains have revealed an age-dependent decline in Nurr1 expression that parallels the reduction of TH-positive dopaminergic neurons in the substantia nigra ^11^. Consistent with these observations, age-associated decreases in Nurr1 expression have been reported in rodent substantia nigra ^12, 13^, as well as in cultured mDA neurons during prolonged in vitro maintenance ^13^. Moreover, Nurr1 heterozygous-deficient mice exhibit accelerated age-dependent loss of nigral dopaminergic neurons compared with wild-type mice ^42, 43^. Together, these findings underscore the essential role of Nurr1 in maintaining mDA neuron integrity and suggest that age-dependent decline in Nurr1 contributes to dopaminergic vulnerability during aging and PD.

JAK2 signaling was originally characterized in immune and hematopoietic systems but has increasingly been recognized as an important modulator of neuronal function. In experimental models of Alzheimer’s disease, JAK2/STAT signaling participates in neuronal survival and stress adaptation in response to amyloid-β (Aβ) toxicity ^23^. Activation of JAK2 via receptors such as the α7 nicotinic acetylcholine receptor, erythropoietin receptor, or leptin receptor attenuates Aβ-induced neurotoxicity through JAK/STAT- or PI3K/Akt-dependent mechanisms ^30, 44^. Similarly, erythropoietin and hydroxy-safflor yellow A (HSYA) protect against Aβ-induced neurotoxicity by modulating the JAK/STAT pathway in PC12 cells ^28^. In vivo studies further support a functional role for JAK2 signaling in hippocampal integrity. For example, intracerebroventricular administration of Aβ suppresses JAK2/STAT3 activity and leads to memory impairment ^29^, indicating that disruption of this pathway compromises neuronal resilience and cognitive function.

Despite these observations, the expression pattern and functional significance of JAK2 in midbrain dopaminergic neurons have not been systematically investigated. In the present study, JAK2 immunoreactivity was minimal in dopaminergic neurons of young adult mice but markedly increased in TH-positive neurons of the substantia nigra in aged mice (12–18 months), indicating age-dependent regulation. Importantly, JAK2 induction was not observed in TH-positive neurons in the 6-OHDA model of acute dopaminergic degeneration, suggesting that JAK2 upregulation represents an aging-associated process rather than a nonspecific response to neurotoxic injury.

JAK2 signaling has also been linked to aging-related processes in several tissues. In the hippocampus, impaired JAK2/STAT3 signaling contributes to cognitive dysfunction in aging models ^45, 46^. Conversely, in certain non-neuronal cells, JAK2 activity has been associated with cellular senescence ^47–49^, highlighting the context-dependent nature of JAK2 signaling. In the present study, our data indicate that JAK2 enhances Nurr1 protein stability and reduces oxidative stress–induced cytotoxicity. These findings suggest that JAK2 functions as an adaptive regulator that helps preserve dopaminergic neuron integrity in the aging midbrain.

Interestingly, our results indicate that JAK2 regulates Nurr1 through a mechanism distinct from canonical JAK/STAT or PI3K/Akt signaling pathways. Instead, JAK2 physically associates with Nurr1 and enhances its protein stability without promoting detectable tyrosine phosphorylation. This phosphorylation-independent mechanism suggests that JAK2 may regulate Nurr1 through protein–protein interactions that influence nuclear receptor stability or turnover. Such a mechanism expands the functional repertoire of JAK2 beyond its classical role as a cytokine-responsive kinase and reveals a previously unrecognized mode of regulation for Nurr1.

A limitation of the present study is that the most robust activation of Nurr1 was observed with the constitutively active JAK2_V617F_ mutant, whereas wild-type JAK2 produced only modest effects in our cellular assays. JAK2_V617F_ has been widely used as a tool to model sustained JAK2 activation, and our findings therefore demonstrate that strong JAK2 signaling is sufficient to enhance Nurr1 stability. Future studies will be required to determine how physiological JAK2 activation regulates Nurr1 in dopaminergic neurons in vivo and whether age-associated signaling pathways contribute to this process. In addition, although our mechanistic analyses were performed primarily in neuronal cell lines, the age-dependent increase of JAK2 expression observed in dopaminergic neurons of the mouse substantia nigra supports the potential physiological relevance of this pathway. Further studies using primary neurons and in vivo models will help clarify the functional role of JAK2–Nurr1 signaling in the aging midbrain.

Previous studies have shown that pharmacological inhibition of JAK1/2 attenuates α-synuclein–induced neuroinflammation by suppressing JAK/STAT signaling in microglia ^50^, highlighting the pro-inflammatory role of JAK signaling in glial cells under pathological conditions. In contrast, our findings suggest that neuronal JAK2 serves a distinct, cell-type–specific function in maintaining dopaminergic neuron integrity during aging. These observations underscore the complex and context-dependent roles of JAK2 signaling in the central nervous system, which warrant further investigation.

In summary, our study identifies JAK2 as an aging-associated regulator of Nurr1 protein stability in midbrain dopaminergic neurons. Given the documented decline of Nurr1 expression during aging, these findings provide new insight into mechanisms underlying age-dependent dopaminergic vulnerability and suggest that modulation of the JAK2–Nurr1 regulatory axis may represent a potential strategy for preserving dopaminergic neuron resilience during aging and neurodegeneration.

## Materials and methods

### Tissue Preparation

Animal care and experimental procedures were conducted in accordance with the guidelines of the McLean Hospital Institutional Animal Care and Use Committee and the National Institutes of Health. Male C57BL/6 mice (3, 6, 12, and 18 months old) were obtained from The Jackson Laboratory (Bar Harbor, ME, USA). Mice were deeply anesthetized and transcardially perfused with phosphate-buffered saline (PBS), followed by 4% paraformaldehyde (PFA) in PBS. Brains were carefully removed and post-fixed in 4% PFA for 24 h at 4°C. Tissues were then cryoprotected by sequential immersion in 10%, 20%, and 30% sucrose solutions until fully equilibrated.

### Partial 6-OHDA lesion model

Animals were anesthetized with isoflurane using a SomnoSuite Anesthesia System (Kent Scientific Corporation, Torrington, CT). Stereotaxic procedures were performed using a stereotaxic apparatus (David KOPF Instruments, Tujunga, CA) equipped with a Micro4 controller (World Precision Instruments, Sarasota, FL). Unilateral lesions targeting the nigrostriatal pathway were induced by the stereotaxic administration of 6-OHDA into the medial forebrain bundle. To protect noradrenergic projections, desipramine (10 mg/kg) was administered 15 min prior to anesthesia. A total of 1 µL of 6-OHDA solution (1.0 mg/mL in 0.2% ascorbic acid and 0.9% saline) was delivered using a 10 µL Hamilton syringe (Hamilton Company, Reno, NV). Injection coordinates were determined relative to bregma as follows: anteroposterior (AP), −1.2 mm; mediolateral (ML), −1.1 mm; and dorsoventral (DV), −5.0 mm. Following surgery, the incisions were closed with Autoclip surgical staples (Fine Science Tools, Foster City, CA), and the animals were maintained on a heating pad for monitoring until they regained consciousness. To alleviate discomfort, all animals received ketoprofen (5 mg/kg, s.c.) and 1 mL of 0.9% sodium chloride (i.p.) for hydration.

### Immunofluorescence

Free-floating midbrain sections were washed three times with PBS and incubated in blocking solution containing 3% bovine serum albumin (BSA) and 0.3% Triton X-100 for 1 h at room temperature. Sections were then incubated overnight at 4°C with primary antibodies against tyrosine hydroxylase (TH; EMD Millipore, MA, USA) or Nurr1 (generated in-house) ^51^. After washing, sections were incubated for 1 h at room temperature with Alexa Fluor 488–conjugated anti-rabbit IgG or Alexa Fluor 546–conjugated anti-mouse IgG secondary antibodies. Fluorescence images were acquired using an Olympus fluorescence microscope (Olympus Corporation, Tokyo, Japan). Relative fluorescence intensity was quantified using ImageJ software (NIH, Bethesda, MD, USA).

### Cell Culture and Transfection

Human neuroblastoma SK-N-BE(2)C cells were maintained in Dulbecco’s Modified Eagle Medium (DMEM) supplemented with 10% fetal bovine serum (FBS) and 100 U/mL penicillin/streptomycin at 37°C in a humidified incubator containing 5% CO₂.

For transfection experiments, cells were plated one day prior to transfection in antibiotic-free DMEM. Transfections were performed using Lipofectamine reagent (Thermo Fisher Scientific, Carlsbad, CA, USA) according to the manufacturer’s instructions.

### Luciferase Reporter Assay

For luciferase assays, SK-N-BE(2)C cells were transfected with reporter plasmids and expression constructs as indicated. For Nurr1-LBD assays, cells were co-transfected with pGAL-Nurr1(LBD), p8xUAS-Luc reporter plasmid, and pRSV-β-galactosidase as an internal control. For full-length Nurr1 assays, cells were transfected with pCMV-Nurr1, p4xNL3-Luc, and pRSV-β-galactosidase plasmids. Cells were additionally transfected with pcDNA3 control vector, JAK2, or JAK2_V617F_ expression plasmids as indicated. Twenty-four hours after transfection, cells were lysed in lysis buffer (25 mM Tris-phosphate, pH 7.8; 2 mM DTT; 2 mM CDTA; 10% glycerol; 1% Triton X-100) for 30 min. Luciferase activity was measured using a luminometer (SpectraMax®, Molecular Devices, Sunnyvale, CA, USA). Transfection efficiency was normalized to β-galactosidase activity.

### Site-Directed Mutagenesis

Point mutations in Nurr1 were generated using the QuikChange II XL site-directed mutagenesis kit (Agilent Technologies, Santa Clara, CA, USA) according to the manufacturer’s instructions. Mutagenesis reactions were performed using pGAL-Nurr1(LBD) or pCMV-Nurr1 templates. All mutations were confirmed by DNA sequencing.

### Quantitative Real-Time PCR

Total RNA was extracted using the GeneJet RNA Purification Kit (Thermo Fisher Scientific, Carlsbad, CA, USA) according to the manufacturer’s instructions. RNA was reverse-transcribed into cDNA using the iScrpit^TM^ cDNA synthesis kit (BIO-RAD, Hercules, CA, USA). For quantitative real-time PCR (qPCR), cDNA samples were mixed with SYBR Green universal master mix (BIO-RAD) and target-specified primers. The primer sequences were as follows:

Mouse JAK2

Forward: 5’-TGCTACGATGCAGCCCTAAG-3’

Reverse: 5’-TGAGCGCACAGTTTCCATCT-3’

Mouse Nurr1

Forward: 5’-CCTGTCAGCACTACGGTGTT-3’

Reverse: 5’-TAAACTGTCCGTGCGAACCA-3’

PCR amplification was performed using a CFX Connect^TM^ Real-Time PCR Detection System (BIO-RAD). Gene expression levels were normalized to GAPDH expression.

### Western Blot Analysis

Total protein lysates were prepared using RIPA buffer (Sigma-Aldrich, St. Louis, MO, USA) supplemented with a protease inhibitor cocktail (Roche, Basel, Switzerland). Protein samples were separated by electrophoresis on Bolt™ 4–12% Bis-Tris Plus gels (Thermo Fisher Scientific) and transferred to PVDF membranes.

Membranes were blocked for 1 h in PBS-T (PBS containing 0.1% Tween 20) supplemented with 5% skim milk. Membranes were then incubated overnight at 4°C with primary antibodies against JAK2 (Cell Signaling Technology, Danvers, MA, USA), c-Myc (Roche Diagnostics GmbH, Mannheim, Germany), or β-actin (Abcam, Cambridge, UK).

After washing, membranes were incubated with HRP-conjugated secondary antibodies for 1 h at room temperature. Immunoreactive bands were visualized using Novex® ECL detection reagent (Thermo Fisher Scientific) and detected using a ChemiDoc™ XRS+ imaging system (Bio-Rad). Band intensities were quantified using ImageJ software and normalized to β-actin.

### Cytotoxicity Assay

Cellular cytotoxicity was assessed by measuring lactate dehydrogenase (LDH) release using an LDH cytotoxicity detection kit (Roche) according to the manufacturer’s protocol. Absorbance was measured at 490 nm using a microplate reader (Synergy HT, BioTek, Winooski, VT, USA).

### Measurement of Oxidative Stress

Intracellular reactive oxygen species (ROS) levels were measured using the fluorescent probe 2′,7′-dichlorofluorescein diacetate (DCFH-DA; Sigma-Aldrich). Cells were incubated with 10 μM DCFH-DA for 30 min at 37°C. After washing with PBS, fluorescence images were acquired using a fluorescence microscope. Fluorescence intensity was quantified using ImageJ software.

### Statistical Analysis

All data are presented as mean ± SEM. Statistical comparisons between two groups were performed using Student’s *t*-test. Multiple group comparisons were analyzed using one-way analysis of variance (ANOVA) followed by Tukey’s post hoc test. Statistical significance was defined as *p* < 0.05 (*), 0.01 (**), or 0.001 (***).

## Supplementary Figures

**Figure S1.** JAK2 expression in the 6-OHDA–lesioned mouse model Immunofluorescence analysis showing that JAK2 immunoreactivity was not detected in degenerating dopaminergic neurons following stereotaxic injection of 6-hydroxydopamine (6-OHDA).

**Figure S2.** Effect of JAK2_V617F_ on Nurr1–RXR**α** heterodimer–mediated transcription RXRα agonists (9-cis retinoic acid and bexarotene) significantly increased Nurr1 transcriptional activity on the DR5 response element. However, JAK2_V617F_ did not further enhance DR5-driven transcriptional activity in SK-N-BE(2)C cells.

**Figure S3.** JAK2_V617F_-induced Nurr1 activation is independent of JAK/STAT signaling (A) Overexpression of STAT3 did not alter JAK2_V617F_-induced Nurr1 transcriptional activation. (B) Pharmacological inhibition of STAT signaling using fludarabine (STAT1 inhibitor), S3I-201 (STAT3 inhibitor), or a STAT5 inhibitor did not suppress JAK2_V617F_-mediated Nurr1 activation.

**Figure S4.** JAK2_V617F_-induced Nurr1 activation is independent of PI3K/Akt signaling Pharmacological inhibition of PI3K (wortmannin and LY294002) or Akt (GSK690693 and MK-2206) did not attenuate JAK2_V617F_-induced Nurr1 transcriptional activation.

**Figure S5.** Prediction of putative JAK2-dependent tyrosine phosphorylation sites in Nurr1 Putative tyrosine phosphorylation sites within Nurr1 were predicted using a group-based phosphorylation prediction system.

