## Supplemental Figure for "Janus kinase 2 regulates Nurr1 protein stability in dopaminergic neurons of the aging midbrain"

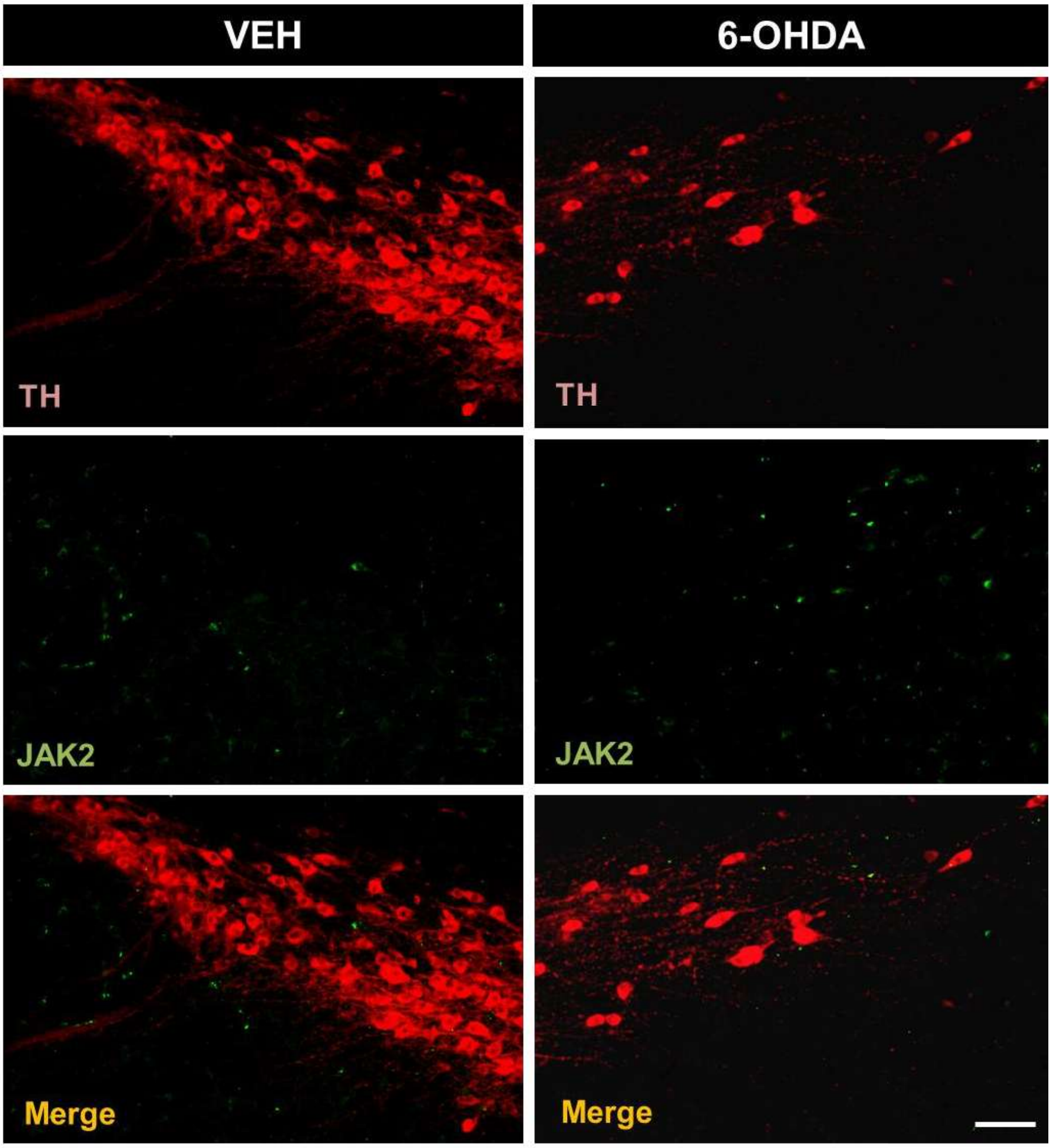


**Figure S1**

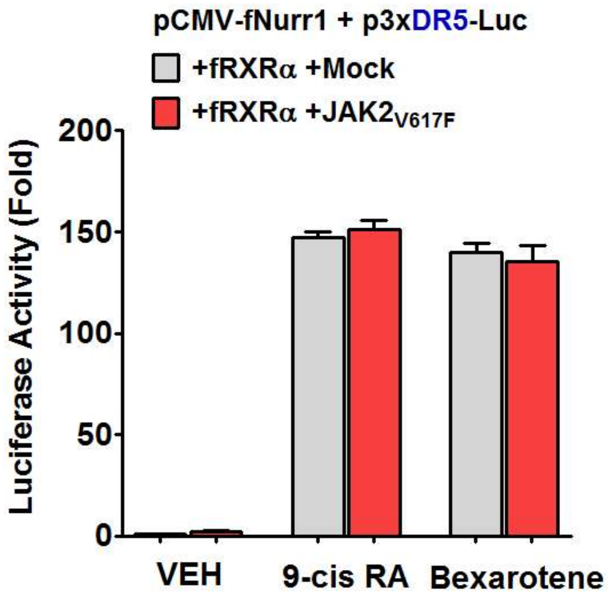
pCMV-fNurr1 + p3xDR5-Luc
+fRXRα +Mock
+fRXRα +JAK2V617F
Luciferase Activity (Fold)
200
150
100
50
0
VEH
9-cis RA
Bexarotene

**Figure S2**

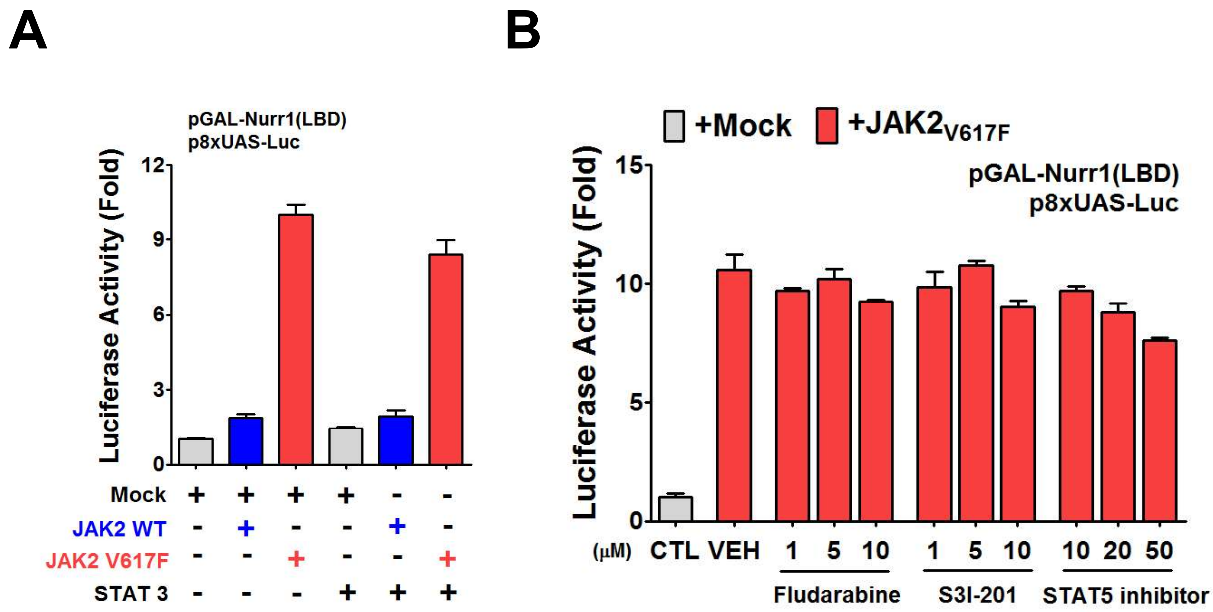
A
pGAL-Nurr1(LBD)
p8xUAS-Luc
Luciferase Activity (Fold)
12
9
6
3
0
Mock + + + + - -
JAK2 WT - + - - + -
JAK2 V617F - - + - - +
STAT 3 - - - + + +
B
+Mock
+JAK2V617F
pGAL-Nurr1(LBD)
p8xUAS-Luc
Luciferase Activity (Fold)
15
10
5
0
(μM) CTL VEH 1 5 10 1 5 10 10 20 50
Fludarabine
S3I-201
STAT5 inhibitor

Figure S3

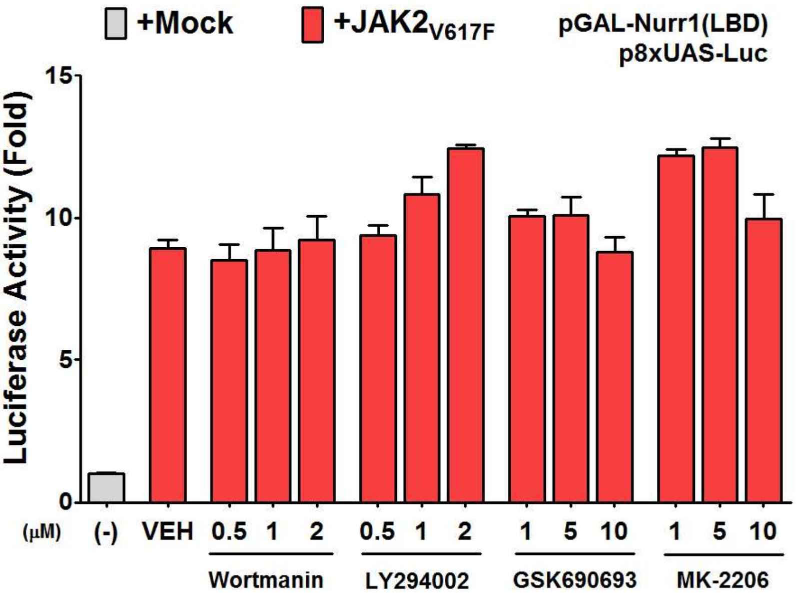

+Mock
+JAK2V617F
pGAL-Nurr1(LBD)
p8xUAS-Luc
Luciferase Activity (Fold)
15
10
5
0
(μM)
(-)
VEH
0.5
1
2
0.5
1
2
1
5
10
1
5
10
Wortmanin
LY294002
GSK690693
MK-2206


**Figure S4**

**Predicted phosphorylation sites by JAKs**

| Position | Code | Kinase | Peptide | Score |
|---|---|---|---|---|
| 384 | Y | TK/Jak/JAK3 | PAMTSLDYSRFQANP | 5.625 |
| 393 | Y | TK/Jak/JAK3 | RFQANPDYQMSGDDT | 5 |
| 575 | Y | TK/Jak/JAK2 | QGLQRIFYLKLEDLV | 10.432 |
| 575 | Y | TK/Jak/JAK3 | QGLQRIFYLKLEDLV | 6.625 |
| 575 | Y | TK/Jak/TYK2 | QGLQRIFYLKLEDLV | 3 |

**Group-based Prediction System**, ver 3.0 (http://gps.biocuckoo.org)

### Figure S5
